# Forecasting bacterial survival-success and adaptive evolution through multi-omics stress-response mapping, network analyses and machine learning

**DOI:** 10.1101/387910

**Authors:** Zeyu Zhu, Defne Surujon, Aidan Pavao, José Bento, Tim van Opijnen

## Abstract

Whether a bacterial pathogen establishes an infection and/or evolves antibiotic resistance depends on successful survival while experiencing stress from for instance the host immune system and/or antibiotics. Predictions on bacterial survival and adaptive outcomes could thus have great prognostic value. However, it is unknown what information is required to enable such predictions. By developing a novel network-based analysis method, a bacterium's phenotypic and transcriptional response can be objectively quantified in temporal 3D-feature space. The resulting trajectories can be interpreted as a degree of coordination, where a focused and coordinated response predicts bacterial survival-success, and a random uncoordinated response predicts survival-failure. These predictions extend to both antibiotic resistance and in vivo infection conditions and are applicable to both Gram-positive and Gram-negative bacteria. Moreover, through experimental evolution we show that the degree of coordination is an adaptive outcome - an uncoordinated response evolves into a coordinated response when a bacterium adapts to its environment. Most surprisingly, it turns out that phenotypic and transcriptional response data, network features and genome plasticity data can be used to train a machine learning model that is able to predict which genes in the genome will adapt under nutrient or antibiotic selection. Importantly, this suggests that deterministic factors help drive adaptation and that evolution is, at least partially, predictable. This work demonstrates that with the right information predictions on bacterial short-term survival and long-term adaptive outcomes are feasible, which underscores that personalized infectious disease diagnostics and treatments are possible, and should be developed.

## INTRODUCTION

The ability to predict whether a bacterial pathogen is successfully establishing an infection, will adapt to the stress it encounters in the host and/or progress to cause disease could have great diagnostic value. However, it is unknown whether such predictions are entirely possible and what information they would require. As a consequence, most diagnostics today come from a physician’s deductive reasoning, which can lead to sub-optimal antibiotic treatments and may contribute to the emergence and spread of antibiotic resistance [1, 2]. Alternatively, in cancer diagnostics transcriptional changes in specific genes of cancerous tissue, in addition to changes in the host response, are used to provide prognostic information beyond standard clinical assessment [3-6]. Moreover, integration of systems-level data, machine learning, and various network/graph-based approaches have been employed to classify cancer subtypes and identify subtype-specific drug targets, enhancing the diagnostic power of current approaches and leading to more effective treatment options [7, 8]. With analogy to cancer diagnostics, a systems-wide understanding of the state of a bacterial infection and how the infection may possibly progress under pressure of the host-immune system and/or other stresses, could similarly aid in providing targeted and personalized infectious-disease treatments.

Our previous work has indicated that advanced infectious-disease prognostics may be possible by combining bacterial stress-response monitoring with network analyses [9]. A commonly applied approach for characterizing bacterial stress responses is through RNA-Seq, which measures genome-wide transcriptional changes upon an environmental perturbation. With the advent of transposon-insertion sequencing (Tn-Seq), it has now also become relatively easy to determine, on a genome-wide scale, the phenotypic importance of a gene, i.e. a gene’s contribution to fitness in a specific environment [10, 11]. Importantly, direct comparisons between data from these different omics-approaches has shown, contrary to expectations, that genes that change in transcription are poor indicators of what matters phenotypically. In other words, phenotypically important and transcriptionally important genes (PIGs and TIGs) rarely overlap [9, 12-18]. However, when integrated into a network, highly coordinated patterns between PIGs and TIGs surface when the organism is challenged with an evolutionarily familiar stress (i.e. one that has been experienced for many generations, e.g. nutrient depletion), while the response becomes less coordinated when the bacterium is challenged with and responds to a relatively new stress (e.g. antibiotics) [9]. This means that the degree of network coordination between PIGs and TIGs originates from the bacterium’s ‘adaptive past’ and should thus be indicative of the degree to which the bacterium is adapted to a specific stress and will survive the challenge (short-term survival outcome). Moreover, since evolution is a continuing process, survival outcome - influenced by past adaptation - is ultimately related to future adaptive outcomes; i.e. network coordination is indicative of where and how stress is experienced in the genome, while selection drives adaptive evolution to resolve this stress. Thus, it may be possible to predict where in the network innovation (adaptation) is most likely to occur to optimize network coordination and increase survival success (long-term adaptive outcome).

Here we develop a novel integrated approach that combines genome-wide profiling, network analyses and machine learning, which enables predictions on bacterial short-term survival and long-term adaptive outcomes. As our model system, we use the respiratory pathogen *Streptococcus pneumoniae*, which on a yearly basis causes ~1 million fatalities worldwide [19] and ~4 million disease episodes in the US alone, among which ~40% are caused by strains that are resistant to at least one antibiotic [20]. To develop this predictive strategy, we first establish the transcriptionally and phenotypically important genes using temporal RNA-Seq and Tn-Seq respectively in different *S. pneumoniae* strains that have different survival outcomes under nutrient stress conditions and in the presence of antibiotics. By overlaying data onto newly developed strain-specific networks and applying network analyses, we find that distinct network patterns emerge that can be depicted as temporal trajectories that move through a specially constructed feature space. Importantly, these patterns are predictive of whether or not a bacterium is successfully surviving in its environment. Moreover, we apply the approach to *in vitro* and *in vivo* data from *Pseudomonas aeruginosa*, highlighting its generalizability and the possibility to predict bacterial survival-success in the host. Lastly, the development of a support vector machine (SVM) leads to the ability to predict which genes acquire adaptive mutations while adapting to nutrient stress or while evolving antibiotic resistance. This study shows that infectious-disease prognostics is feasible through the implementation of different omics-approaches, network analyses and machine learning, enabling the prediction of whether a bacterium will survive or not under a given stress and where in the genome it is most likely to adapt.

## RESULTS AND DISCUSSION

### Strain specific metabolic networks are insufficient in defining nutrient dependency or predicting survival outcomes in three strains of *S. pneumoniae*

*Streptococcus pneumoniae* on average contains 2100 genes and harbors considerable genetic diversity, with two strains differing on average by 250 genes (presence and absence), and a pan-genome (collection of all genes across all strains) that is approximately double the size of the genome of any given strain. *S. pneumoniae* designates ~30% of its genome to metabolic functions, which enables growth on different carbon sources and in the presence and absence of different substrates (e.g. amino acids, lipids). This ‘strategy’ all but guarantees the bacterium’s survival in a variety of host-niches, including the nasopharynx, inner-ear and lungs. Since different host niches have different nutrient availability [21], nutrient depletion is evolutionarily an important stress to the obligate non-motile human pathogen *S. pneumoniae* and has shaped its genetic composition. We thus reasoned that strain-specific nutrient dependencies must exist and that such dependencies can be used as a testing-ground to predict whether a strain will survive in a specific environment and what information is needed to make such predictions.

Three strains (TIGR4 [T4], Taiwan-19F [19F] and D39) that differ in ~7% of their genetic content (presence or absence of genes; [9]), were assayed to identify essential nutrients for growth. Single nutrients were sequentially removed from a chemically defined medium (CDM) and the effect on the growth rate was calculated. A nutrient is defined as essential if its removal causes a >70% reduction in the bacterium’s growth rate, and important if the reduction is between 50-70% (Supplementary Figure 1, detailed explanation of definitions in this study can be found in Supplementary Information). In total, four amino acids are essential to all three strains: (L-Arginine, L-Cysteine, L-Histidine, and L-Leucine; Supplementary Figure 1A), while 6 nutrients have strain-specific requirements: 1) three amino acids (Glycine, L-Isoleucine and L-Valine) and the nucleobase uracil are essential to D39; 2) Pantothenate is important to T4; 3) L-Glutamine is important for T4 and D39 (Supplementary Figure 1A). At least two possible explanations for this strain-specific nutrient dependency are that a strain either lacks certain genes that are required to synthesize the nutrient or the respective metabolic network is differentially wired. For instance, a metabolic gene might encode isoforms of an enzyme that catalyze different reactions in different strains [22]. To determine the origin of the strain-specific nutrient dependency we expanded the *S. pneumoniae* metabolic model we previously built for T4 [9] with two additional strain-specific models for D39 and 19F (Supplementary Figure 2; Supplementary File 2). The three models are highly conserved, sharing 96% of all metabolic genes across all strains (i.e. 431 metabolic genes/868 metabolic reactions), however, neither the presence of strain-specific metabolism genes nor differences in the metabolic network topology can sufficiently explain the observed strain-specific nutrient requirements.

### Genome-wide profiling reveals distinct transcriptional patterns between a nutrient dependent strain and an independent strain

Genomic content and network architecture are thus not enough to consistently predict bacterial survival and growth in a certain environment. We previously demonstrated that the degree of network coordination between phenotypic and transcriptional responses distinguishes evolutionarily familiar stresses from relatively novel ones [9]. Such network patterns could thus be key to predicting whether a bacterium is successfully surviving in a specific environment.

To uncover genes that are phenotypically important (PIGs), we performed Tn-Seq on T4 in the absence of either uracil, L-Valine or Glycine (i.e. nutrients essential for D39 but not T4). Tn-Seq measures, in a highly quantitative fashion and on a genome-wide scale, which genes and pathways are important for growth in a specific environment [11, 23]. By comparing fitness in the presence and absence of a nutrient, genes that are important for T4’s survival in the absence of the nutrient are identified, which leads to a total of 134 PIGs that contribute to growth of T4 (15 genes for Glycine, 75 genes for uracil, 44 genes for L-Valine). All of these genes have homologs in D39 and thus do not directly explain the different dependencies between T4 and D39. Subsequently, we profiled the manner in which T4 and D39 transcriptionally respond to the absence of the D39-specific essential nutrients. Genome-wide transcriptional responses were determined by temporal RNA-Seq for T4 (the nutrient-independent strain) and D39 (the nutrient-dependent strain) at 30 and 90 min after nutrient depletion (Supplementary Table 1).

Three distinct transcriptional patterns emerge that differentiate a nutrient-dependent from an independent strain: 1) A dependent strain tends to trigger a greater number of expression changes under nutrient depletion (Supplementary Table 2). For instance, in the absence of L-Valine or Glycine, D39 triggers significantly more TIGs than T4 at both the early and the late time points (two proportion Z-test, p<0.01) (Supplementary Table 2). Additionally, in the absence of uracil, D39 and T4 trigger similar numbers of TIGs at 30min, however at 90min, the number of TIGs in T4 decrease (from 22 to 13), while in D39 the number of differentially expressed genes increases to 857 (nearly 40% of the genome); 2) In each single nutrient-depletion condition, magnitude distributions of differential expression are significantly wider in D39 than in T4 (Figure 2A, Kolmogorov-Smirnov test, p<0.01, Supplementary Table 2), indicating that the extent of genome-wide transcriptional change is much larger in the dependent strain; 3) A functional distribution analysis of TIGs shows that at 30 and 90 min after the depletion of Glycine or L-Valine, and at 90 min after the depletion of uracil more TIGs per functional tag are differentially regulated in the dependent strain (Figure 2B; Supplementary File 3). Furthermore, the TIGs are distributed across more functional categories indicating that nutrient depletion has a greater impact on most cellular systems of the dependent strain (Figure 2C; Supplementary File 3). If we directly compare the TIGs of the independent with the dependent strain, it turns out that the T4-TIGs (both early and late) are also TIGs in D39. This suggests that the dependent strain can raise a similar ‘appropriate response’ as the independent strain to the endured stress. To obtain slightly higher temporal resolution we additionally profiled 60 min after uracil depletion, which triggers 20 TIGs in D39, the majority of which are involved in uracil uptake (uracil permease SP_1286) and the metabolic pathway that generates the pyrimidine precursor uridine monophosphate (UMP) (SP_0701-0702, SP_0963-0964, SP_1275-1278, SP_1288; Supplementary File 3). These exact uracil-related genes are also up-regulated in T4 and form the majority of T4’s response to uracil depletion at both early and late time points (Supplementary File 3, Figure 2D). Importantly, this further shows that D39 is actually able to generate an appropriate transcriptional response, but only over a limited amount of time. Instead, somewhere between 60 and 90 minutes D39’s response is washed out by a rapidly expanding genome-wide dysregulation (Figure 2E-F).

**Figure 1.**
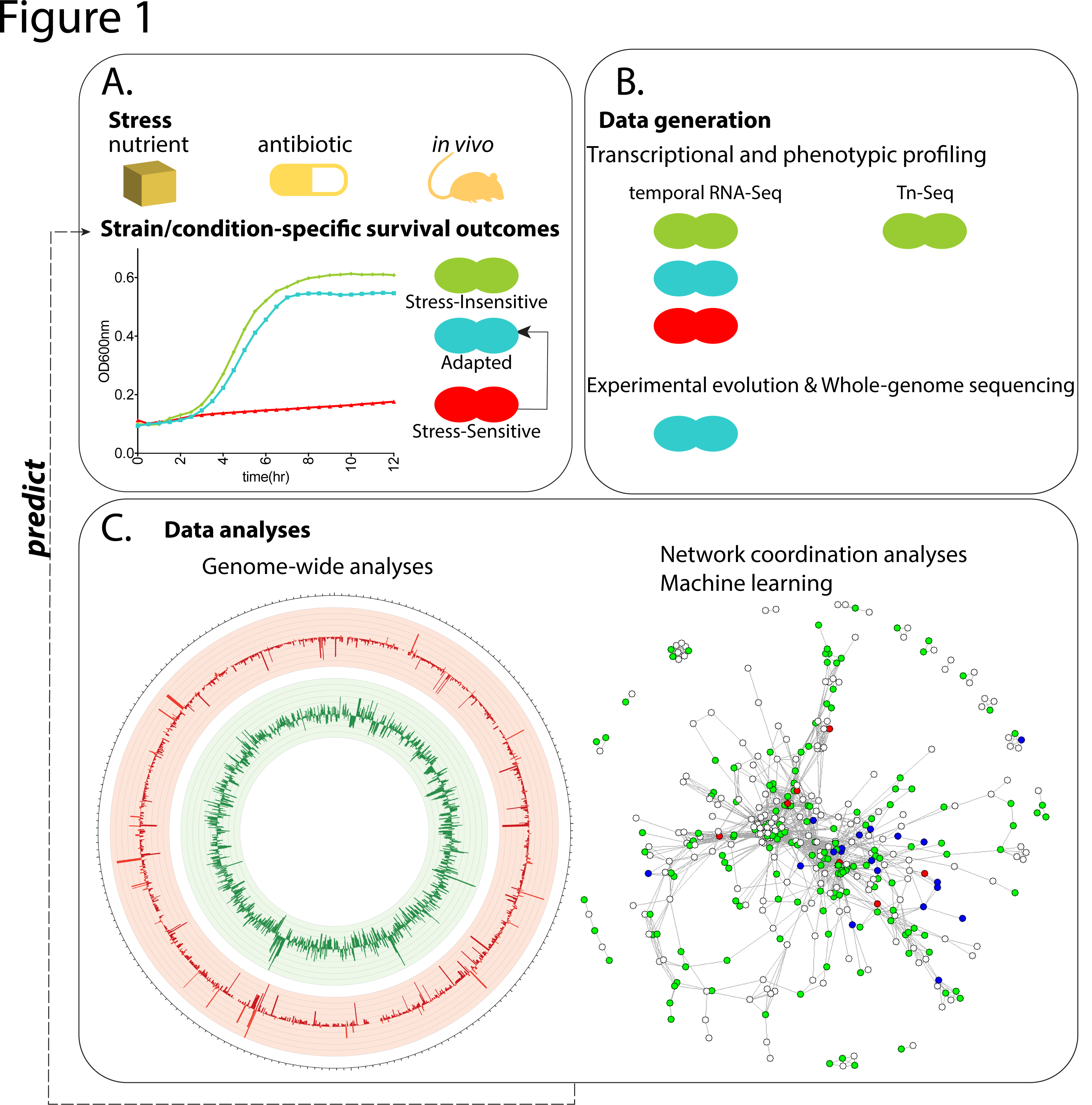
Study overview. **A.** Differential survival outcomes under nutrient depletion, antibiotic exposure and *in vivo* conditions from *Streptococcus pneumoniae* and *Pseudomonas aeruginosa* are investigated in this study. Experimental evolution is performed on stress-sensitive strains (red) to achieve adapted strains (blue). **B.** Temporal RNA-Seq data are collected from the stress-insensitive (green), stress-sensitive and adapted *S. pneumoniae* strains; Tn-Seq data are collected from the stress-insensitive strain. RNA-Seq and Tn-Seq data of *P. aeruginosa* are obtained from published datasets (Murray *et al*, 2015, Turner *et al.*, 2014). **C.** Data obtained from (**B.**) are subjected to genome-wide analyses, network coordination analyses and machine learning to generate predictive patterns of survival outcomes for the stress-sensitive, insensitive and adapted strains; and adaptive outcomes for the stress-sensitive strains.

**Figure 2.**
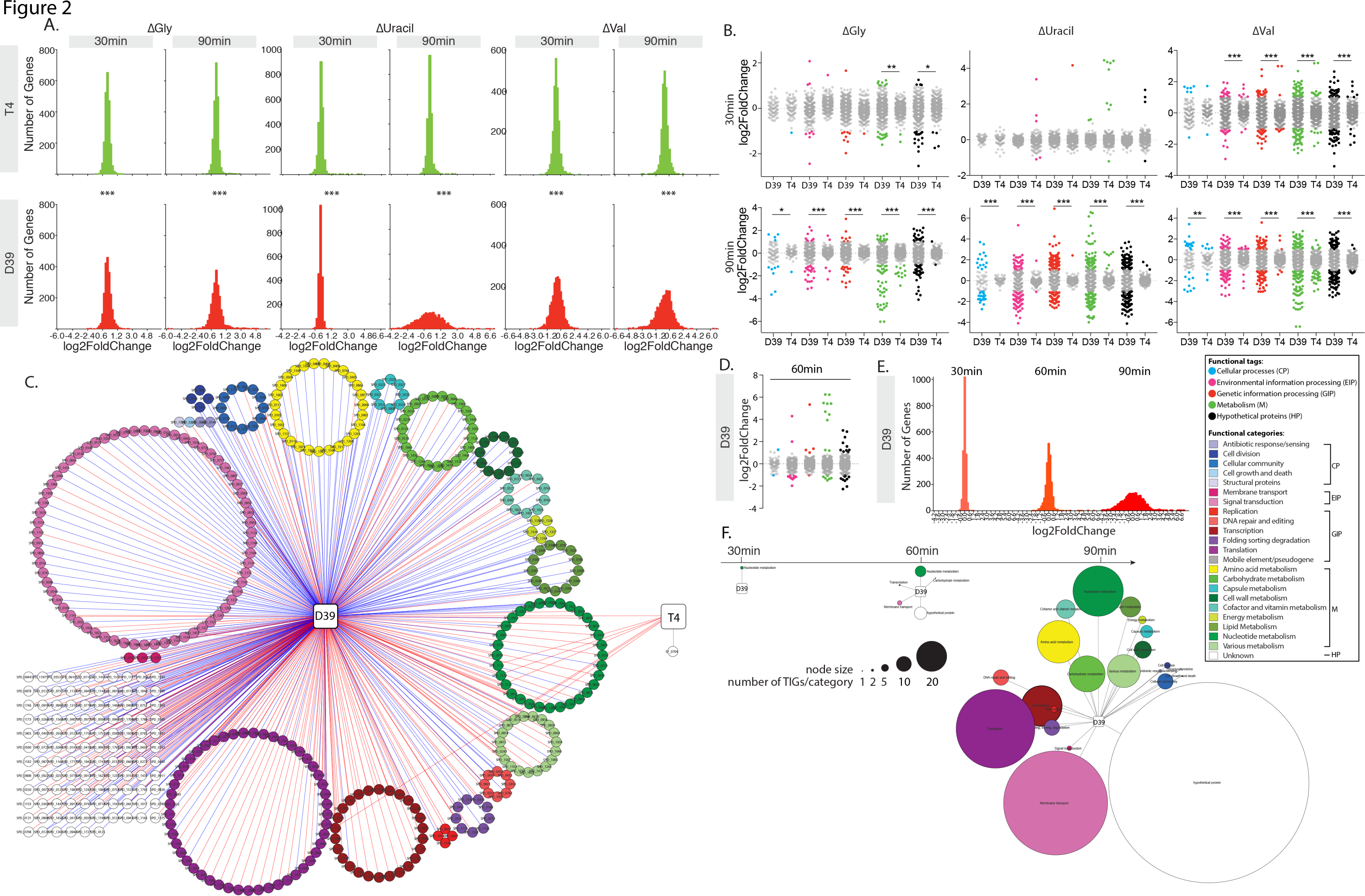
Distinct patterns characterize the transcriptional response of nutrient-dependent and nutrient-independent strains. **A.** The magnitude of genome-wide differential expression shows significantly different distributions between D39 (red) and T4 (green) in the absence of Glycine (ΔGly), uracil (ΔUracil) or L-Valine (L-Val) at 30min and 90min in a Kolmogorov-Smirnov test. **B.** D39 triggers significantly more TIGs in each functional tag than T4, compared in a Z test for two population proportions with Bonferroni correction for multiple testing. **C.** Genome-wide functional category distribution of TIGs in D39 and T4 after 90 minutes of uracil depletion. D. Functional tag distribution of TIGs in D39 after 60 minutes of uracil depletion resembles T4. Genome-wide differential expression of D39 under uracil depletion shows time-dependent increase in magnitude (**E.**) and function distribution (**F.**). For **A-B**, *: 0.001<p<0.02; **: 0.0001<p<0.001; ***: p<0.0001. See in-figure legend for color-coding schemes of functional tags and categories in B- D, F.

### Network analyses of the transcriptional and phenotypic responses can be visualized in a temporal feature space and define survival as a coordinated response

To enable detailed network analyses and determine the degree of network coordination, the strain-specific metabolic network models were converted into genetic networks where each gene is represented as a node, and two gene nodes are connected if the proteins encoded by these genes are involved in the same or in subsequent reactions. Overlaying TIGs and PIGs on the network shows very little overlap and when genome-wide fitness is plotted against expression change most genes distribute along the horizontal and vertical axes (Supplementary Figure 3). This means that genes that change in expression rarely change in fitness, indicating that transcriptional importance is a poor indicator of what matters phenotypically (Supplementary Table 2, Supplementary Figure 3), which is consistent with our previous observations [9]. When the independent and dependent strains’ responses are plotted on a network, visual inspection suggests that the independent response remains contained to a specific part of the network over time (Figure 3A), while the dependent strain’s response becomes increasingly scattered across the entirety of the network (Figure 3B). In order to objectively quantify these responses we devised three types of measurements that capture the defining network characteristics of a response:

1. Connectedness (CC): the number of connected components is calculated by removing all nodes from the network that are neither PIGs nor TIGs. This leaves a collection of sub-networks (or components) that are separated and unreachable from one another. In a network sense, this means that information may flow within a component but not between components due to missing connections. The number of components thus explains the cohesiveness and continuity of the response. For instance, in the absence of uracil in T4 we observe one large component which corresponds to the UMP biosynthesis pathway, and several small (single-node) components (Figure 3C). In contrast, the dependent D39-uracil at 90 min response is defined by a large number of small components consisting of 1 or 2 genes (Figure 3D), however a large dominating component consisting of 121 TIGs and PIGs is also observed (Figure 3D). This large component potentially results from the presence of few highly connected “hub” genes. It is thus important to evaluate whether the number of connected components formed in an observed response are significantly different from a random response, which is achieved by permutation testing (see Methods).
2. Closeness (CN): while a small number of components may indicate that a response is contained to a few network modules, it is equally important to take into account the relative position, or closeness, of the components, where highly related (sub)pathways are generally closer to each other than unrelated pathways. This measure thus explains whether components are functionally related and a response is targeted. For instance, out of the 13 components in the T4 uracil depletion response 12 are only 2-3 edges away from their nearest component (Figure 3E), which is significantly smaller than the distances between randomized responses (obtained through permutation testing). This indicates that the response, while not fully connected, is contained and targeted in a relatively small area of the full metabolic network. In contrast, the distances between the components of the D39-uracil response are not significantly smaller than a random response (Figure 3F).
3. Representation (RE): while our network is limited to metabolism, the observed TIGs and PIGs are genome-wide (Figure 2). For instance, D39’s response to Glycine depletion is significantly connected, however the metabolic portion of the response comprises only ~20% of the full response. Importantly, since only the part of the response that falls on the network is considered, the majority of the response in this case is thus ignored. This heavily skewed off-network response is problematic because while the 20% on-network may give an indication of being connected and/or close, in reality the true response could be random. This is illustrated with respect to the earlier observation that even though the dependent strain may trigger an appropriate transcriptional response that suggests survival-success, when the entire response is considered it becomes clear that the transcriptional dysregulation is scattered across many other non-metabolic pathways, processes and genes that are overwhelming the “appropriate response” (Figure 2). To account for this, the RE is calculated, which defines a response as “metabolically represented” if a significant proportion of the responsive genes fall on the metabolic network (see Methods).

**Figure 3.**
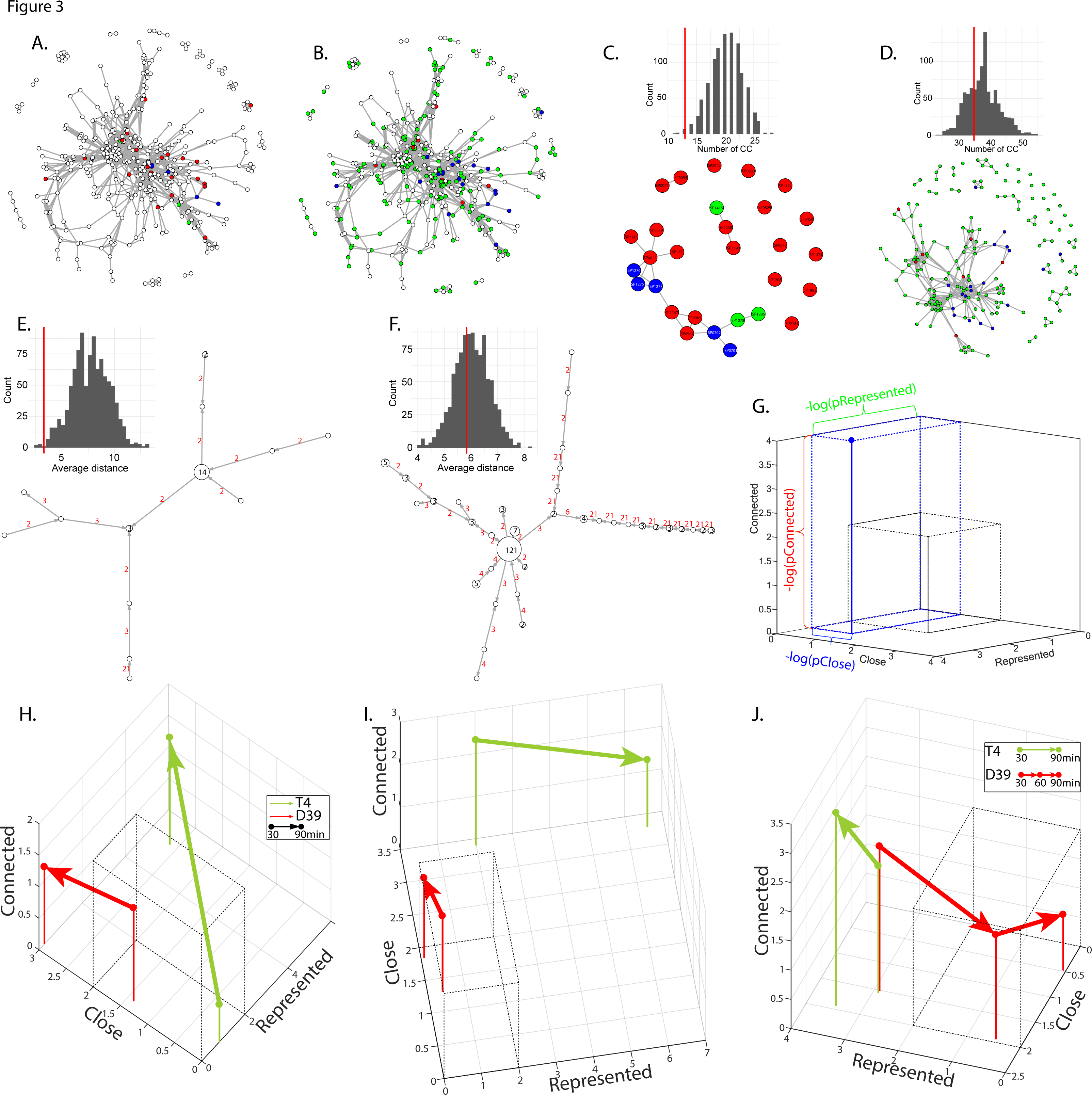
Network coordination analyses can be visualized in a feature space and define survival as a coordinated response. PIGs (red), TIGs (green) and PIG/TIG overlaps (blue) from the uracil depletion experiment (at 90 minutes) are overlaid on the metabolic network for TIGR4 (**A.**) and D39 (**B.**), highlighting differences in network response. Connected components (CC) formed by PIGs and TIGs and the shortest path distances between CC are calculated for TIGR4 (**C.** and **E.**) and D39 (**D.** and **F.**).**C-F**. Inset histograms show the expected results (permutation testing) in comparison with experimental observations (red lines). The p-value is the proportion of permutations that are more extreme than the observation. **G.** Example of the integration of the three-coordination metrics (CC, CN, and RE) for an experiment (blue point) by plotting the -log(p-value) in a 3-dimensional feature space. The gray box represents the significance threshold for each p-value. A coordinated response is typically far away from the origin.**H-J.** Response trajectories for D39 (red) and T4 (green) from 30 to 90 minutes in the absence of L-Valine, Glycine or uracil, respectively. In (**J.**) the D39 trajectory includes the 60- minute time point. For all three graphs the dependent strain D39 remains close to the origin (uncoordinated response), while the independent strain T4 moves away from the origin (coordinated response). An alternative visualization of the degree of coordination of each individual data point can be found in Supplementary Figure 4.

Lastly, to incorporate the manner in which the response changes over-time the log-transformed p-values for CC, CN and RE calculated from each time point are plotted in a feature space, where each of the three characteristics are placed along separate axes (Figure 3G; Supplementary Figure 4). In this scheme, the region around the origin (grey box, Figure 3G) represents a response that is non-significant in terms of CC, CN, and RE.

For all three depletion conditions (L-Valine, Glycine and uracil), the response of the nutrient-independent strain (T4) tends to move away from the origin over time, and the responses are characterized by significant CC, CN and/or RE (Figure 3H-J). In contrast, the nutrient-dependent D39’s responses are mostly confined to the non-significant regions near the origin (Figure 3H-J).
This is especially well illustrated by the uracil depletion experiment, where T4 and D39 strains are situated at a very similar location at 30 min (Figure 3J). However, while the independent strain T4 moves towards a higher CC, CN and RE, D39 moves in the opposite direction and into the non-significant space. Thus, coordination between the transcriptional and phenotypic response is maintained and strengthened over time in strains that can tolerate and survive in a particular environment but weakened in strains that cannot (Figure 3H-J). Importantly, this trajectory reinforces quantitatively what was suggested by the transcriptional response where both strains start out in a very similar manner, and while the T4 response remains targeted, the D39 response ends in uncoordinated dysregulation (Figure 2-uracil depletion). The temporal trajectory formed by three network parameters (CC, CN, RE) thus characterizes the stress-response of a strain as coordinated or uncoordinated, corresponding to survival success or failure.

### Experimental evolution of a sensitive strain reverts nutrient dependencies and rewires stress responses into a coordinated response

In order to test whether a dependent strain that becomes adapted to the absence of a nutrient (i.e. it becomes independent) acquires network coordination, two short-term evolution experiments were designed in which D39 was adapted to grow in the absence of uracil or L-Valine separately. Four replicate populations were established for each experiment and cultured by serial passaging in CDM in which either nutrient was decreased by approximately 15% every 3 days until populations were obtained that are able to robustly grow in the absence of either nutrient (~40 generations each; Supplementary Figure 1C; Figure 4A).

**Figure 4.**
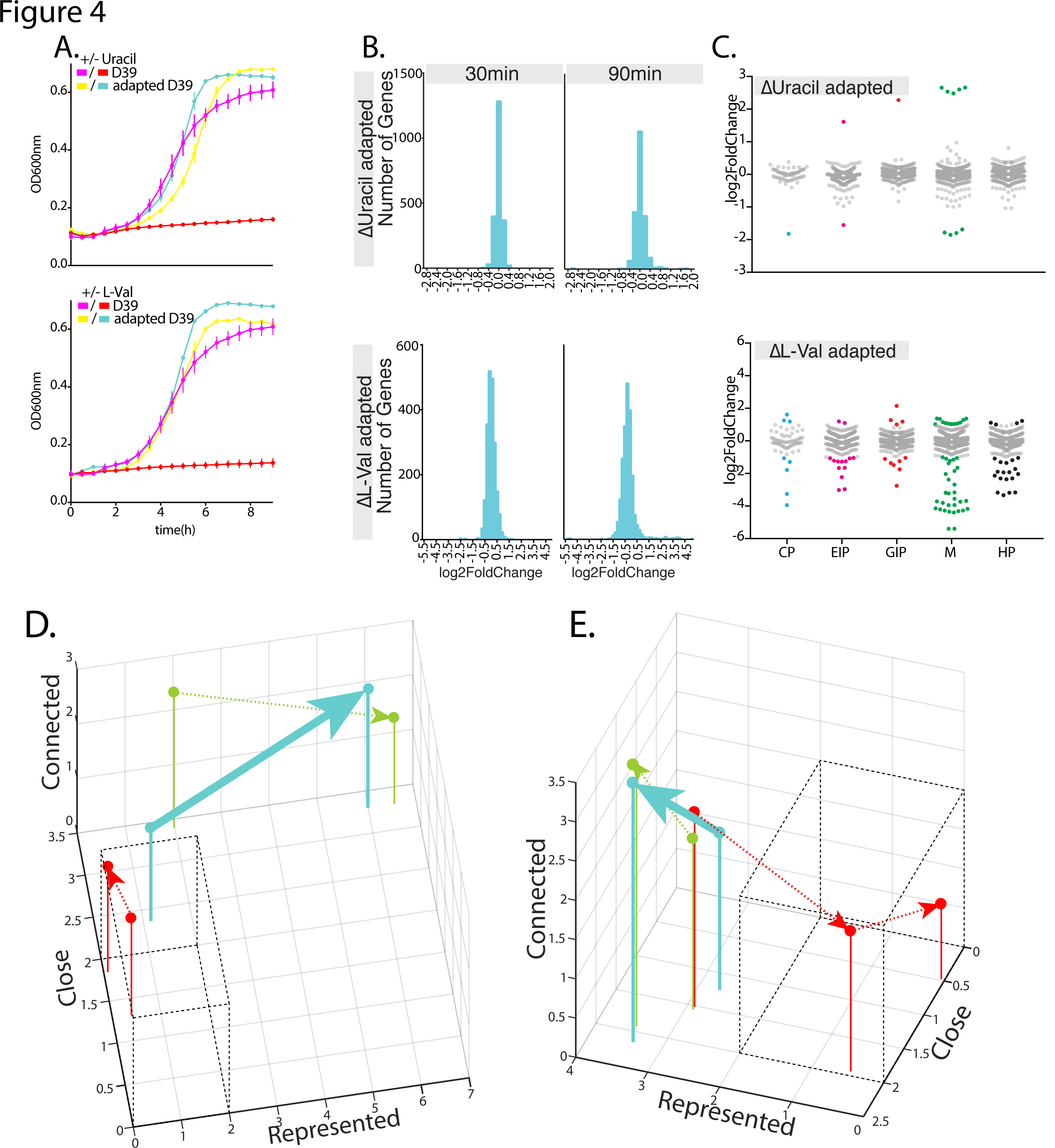
Experimental evolution revert nutrient dependencies and rewires stress responses into a coordinated response. **A.** Adapted D39 strains recover growth in the absence of uracil (top; aD39-uracil) or L-Valine (bottom; aD39-val). **B.** Differential expression magnitude distributions are narrower in aD39-uracil and aD39-val compared to D39and resemble T4 (Figure 2A). C. Functional tag distribution of TIGs in aD39-uracil and aD39-val at 90min after uracil or L-Valine depletion are narrower compared to D39 and resemble T4 (Figure 2B). Network trajectories of aD39-uracil (**D.** blue) and aD39-val (**E.** blue) show an increase in coordination from 30 to 90 minutes that are similar to T4 (**D.** and **E.** green) and dissimilar to wild-type D39 (**D.** and **E.** red).

To determine the adapted strains’ transcriptional response, temporal RNA-Seq was performed on a uracil-adapted (aD39-uracil) and a L-Valine-adapted strain (aD39-val) in the presence and absence of the respective nutrient. Similar to the original independent strain T4, the two adapted D39 strains now exhibit only a small number of differentially expressed genes (Supplementary Table 2; Supplementary File 3), the magnitude of differential expression has a narrow distribution, and TIGs in the adapted strains show specific function distributions similar to the ‘original’ independent strain T4 (Figure 4C and Figure 2B). On a network level, coordination profiles and trajectories arise that are highly similar to T4 (Figure 4D and E). For instance, the trajectory of aD39-val tracks along a higher RE and CC, resembling T4 (Figure 4D), and the trajectory of aD39-uracil moves in the opposite direction of D39 with higher CC, CN and RE and is almost indistinguishable from T4 (Figure 4E). Our analyses thus show that adaptation to nutrient depletion stress leads to transcriptional rewiring and that adapted strains gain highly targeted and coordinated responses, predictive of their ability to survive in an environment.

### Rewiring of genome-wide transcriptional and phenotypic responses to achieve coordination extends to the evolution of antibiotic resistance

To test if network trajectories can also predict survival outcomes in a more complex environment, we extended our approach to the evolution of antibiotic-resistance by challenging T4 with vancomycin. Vancomycin is often used in treating infections caused by beta lactam-resistant *S. pneumoniae* especially during sepsis and meningitis [24, 25]. The MIC of T4 is 0.24ug/mL (Supplementary Figure 1C) and in order to obtain a vancomycin-adapted strain a short-term evolution experiment was performed. Four replicate populations were adapted to vancomycin for ~70 generations (Supplementary Figure 1C), and an adapted strain (aT4-vanc) was isolated, which can grow at 1xMIC with a relative fitness of *W_aT4-vanc_* = 0.88 compared to the no drug control (Figure 5A; i.e. a 12% relative growth defect). Fluorescence microscopy on T4 (wild-type) and aT4-vanc reveal significantly longer cell chains for T4 in the absence of vancomycin (p<0.0001 in t-test; Figure 5B and C). After one-hour exposure to vancomycin (1xMIC), the wildtype loses the long chain morphology and often exhibits a bulging phenotype (Figure 5B), which is in agreement with previous reports [26], while aT4-vanc cells under vancomycin treatment are indistinguishable from untreated cells (Two-sample t-test, p=0.6001) confirming their adapted state.

**Figure 5.**
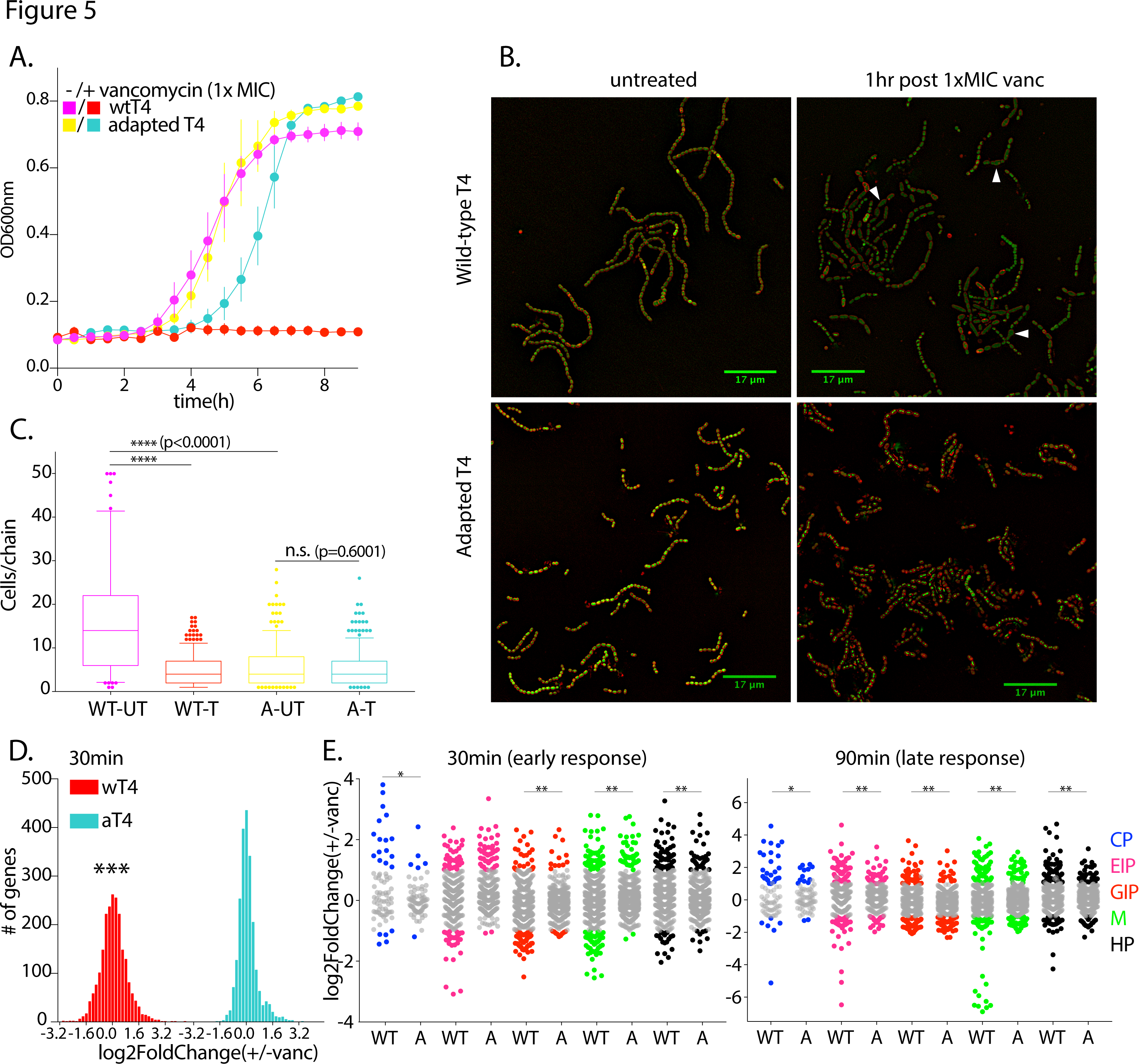
Adapted *S. pneumoniae* exhibits reduced sensitivity, changed morphology and a rewired transcriptional response under vancomycin treatment. Growth phenotypes (**A.**) and morphology (**B.**) of wild-type (WT) and adapted T4 were examined in the absence and presence of vancomycin (1xMIC) in SDMM. **B.** Cells were stained with Syto9 (green) and fm464 (red). White arrowheads highlight bulging cells, typical of vancomycin sensitivity. **C.** Cell numbers per chain were quantified from 1000 cell chains, indicating the adapted strain has a shorter chain-length phenotype, comparable to the vancomycin-treated WT. **D.** Genome-wide differential expression shows a significantly wider magnitude distribution in WT compared to adapted T4 at 30min post-vancomycin treatment in a Kolmogorov-Smirnov test. **E.** WT triggers significantly more TIGs than adapted T4 in most functional tags in both early and late vancomycin response in a Z-test for two population proportions with Bonferroni correction for multiple testing in (**E.**). n.s.: p>0.02, *:0.001< p<0.02; **:0.0001<p<0.001, ***: p<0.0001

The transcriptional response of T4 and aT4-vanc was determined with RNA-Seq at six time points post-vancomycin treatment (10, 20, 30, 45, 60, and 90 min at 1xMIC). Overall, the distinct patterns that are observed under nutrient-depletion are observed in the presence of vancomycin as well: 1) aT4-vanc triggers fewer differential expression than T4 (Supplementary Table 2); 2) aT4-vanc has significantly narrower magnitude distributions of differential expression (Figure 5D, Supplementary Table 2; Kolmogorov-Smirnov test, p<0.02); 3) aT4-vanc triggers significantly fewer TIGs in most functional tags (Figure 5E and Supplementary Table 2; p<0.002 with Bonferroni correction for multiple testing).

To generate the phenotypic response and enable network analyses Tn-Seq was performed in the presence of vancomycin, which, as expected, reveals little overlap between PIGs and TIGs (Supplementary Table 2). The CC, CN and RE trajectories for T4 and aT4-vanc at 1xMIC start at very similar coordinates in the feature space with high RE (Figure 6A). However, T4 rapidly transitions to a less-represented space,displaying an erratic trajectory that ends in a non-significant and uncoordinated response, indicative of survival-failure. On the other hand, aT4- vanc moves away from the origin, to a state where it is significantly connected, close and represented over the first 30 minutes. Between 30 and 90 minutes, aT4-vanc then follows an arc where it gradually becomes less represented, close or connected, and eventually ends just below the significance threshold for all three characteristics (Figure 6A). Thus, while aT4-vanc can maintain a highly coordinated response for at least 60 minutes, this coordination is still partially lost at the 90-minute time point, most likely because aT4-vanc is not fully adapted to vancomycin, displaying a detectable growth defect in the presence of 1xMIC compared to the no drug control (Figure 5A). We reasoned that at a higher vancomycin concentration, aT4-vanc would start to behave more similarly to the sensitive T4 at 1xMIC. When challenged with 1.4xMIC of vancomycin, aT4-vanc initially shows a similar trajectory to 1xMIC (Figure 6A) but traverses the same arc faster, i.e. at 1xMIC aT4-vanc traverses an arc over 60 minutes whereas at 1.4xMIC the traversal of the same arc is completed in 30-45 minutes. Finally, at 1.4xMIC, between 45 and 90 minutes, the trajectory stays near the non-significant space. Thus, aT4-vanc at 1.4xMIC displays similarities to both the wild-type and aT4-vanc at 1xMIC where it has a coordinated response at earlier time points but loses its coordination relatively fast (over fewer number of line segments) and behaves erratically (similar to T4) at the later time points. This means that, similar to nutrient-depletion, the direction of the trajectory but also the shape and the speed at which it moves along a trajectory has predictive value concerning short-term survival success under antibiotic exposure.

**Figure 6.**
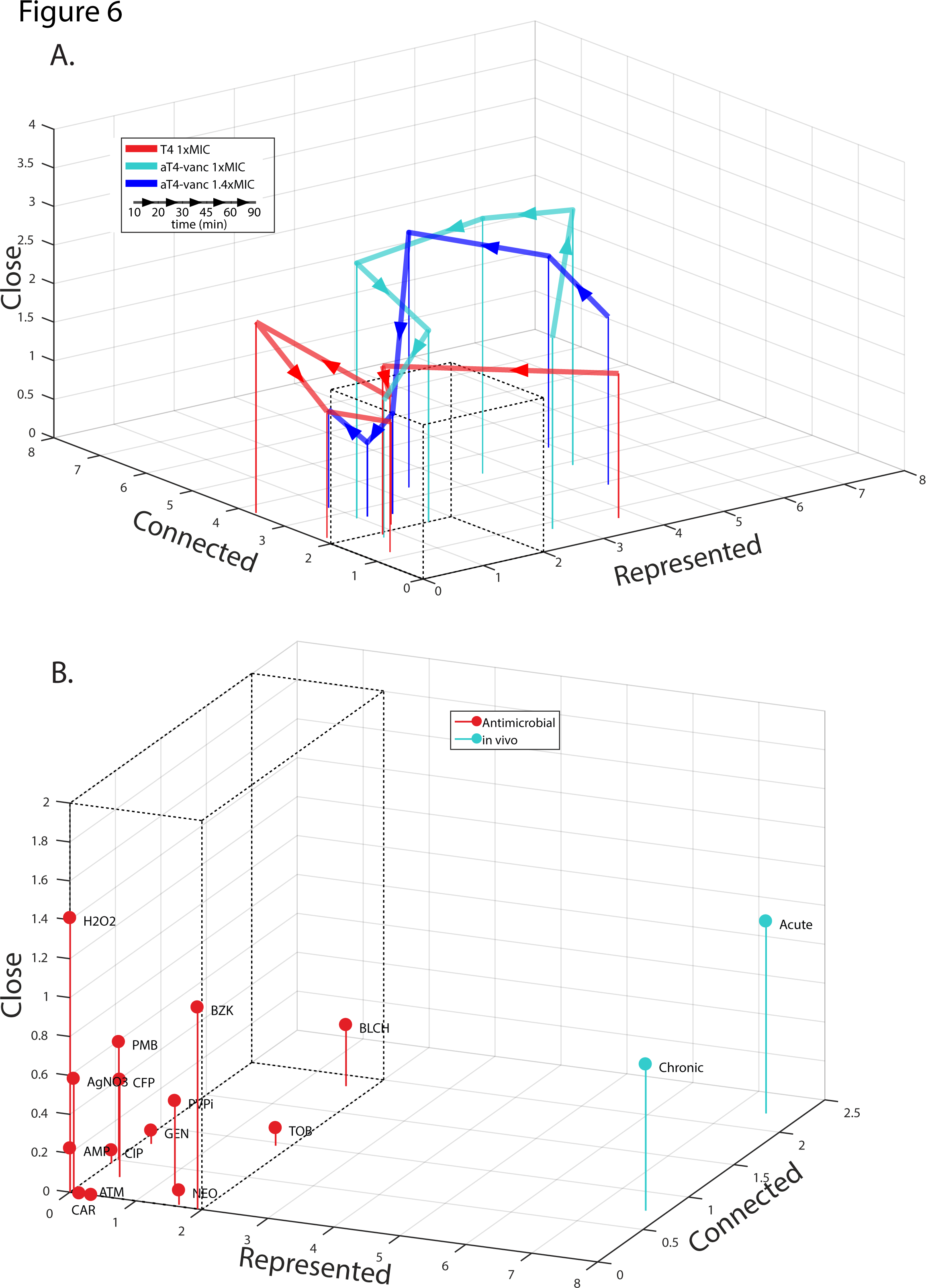
Network coordination defines antibiotic resistance in S. *pneumoniae* and antimicrobial and *in vivo* responses in *P. aeruginosa*. **A.** Temporal network trajectories of the vancomycin response for vancomycin-sensitive (wild-type T4, blue) and vancomycin-adapted (aT4-vanc, red) strains profiled at 10, 20, 30, 45, 60 and 90 minutes after 1xMIC vancomycin treatment. In addition, aT4-vanc is also profiled under 1.4xMIC vancomycin (green). All three trajectories start at a significantly represented state, however the T4 response quickly becomes uncoordinated and erratic. In contrast, aT4-vanc demonstrates a gradual trajectory that mainly moves through significantly coordinated intermediate time points. N.B the speed at which a trajectory is traversed is determined by the number of line segments, and not by the lengths of segments, as each line is a separate time point. **B.** Network coordination analyses extended to *P. aeruginosa* distinguishes between uncoordinated responses to antimicrobials (red), and coordinated responses in *in vivo* wound infection models (blue).

### Network coordination is predictive of survival outcome in other bacterial pathogens

In order to determine whether our findings are applicable to other bacterial species, network analyses were extended to the evolutionarily distant opportunistic pathogen *Pseudomonas aeruginosa* [27]. Tn-Seq and RNA-Seq data collected for strain PAO1 tested against 14 antimicrobials [28] and for strain PA14 tested in 2 *in vivo* wound infections (chronic and acute) [17] were overlaid onto their respective strain-specific metabolic models [29]. In none of the 14 antimicrobial conditions PA14 elicits a coordinated response, i.e. CC, CN and RE of the PIGs and TIGs are never significant (Figure 6B). On the other hand, during an infection, the transcriptional and phenotypic responses of PAO1 are significant in RE on the metabolic network, and in the case of an acute infection the response is also significant in CN (Figure 6B). The higher coordination in the acute infection suggests that the pathogen is more likely to survive in this condition. Indeed, acute burn infections tend to spread and deteriorate rapidly [30], indicating a more successful outcome (at least with respect to short-term bacterial survival) for the pathogen *P. aeruginosa*, and thus suggesting that network analyses can be applied to infer disease progression, although more time-points would most likely be more informative.

### Integration of machine learning, genome-wide profiles, and network characteristics enables prediction of adaptive evolution

Network analyses thus reveal where on the genetic network stress is experienced, while the level of coordination is indicative of how stress is processed. Importantly, adaptive mutations are generally localized in genetic regions that resolve (part of) the experienced stress. It may thus be possible, that with the right information (e.g. where is stress experienced in the genome, how evolvable is that part of the genome, how is it connected in a network context), we can predict which parts of the genome are most likely to contribute to adaptive evolution. Since there are no obvious patterns in our data (e.g. TIGs, PIGs, network connectivity) that are predictive of adaptation we test this hypothesis by training a support vector machine (SVM) - one of the most established supervised classifiers in machine learning [31], with the goal to develop a model that is able to predict which genes will acquire adaptive mutations.

Adaptive mutations are defined as non-synonymous mutations in coding regions that went to fixation or reached a frequency > 50% during experimental evolution in the absence of uracil and L-Valine and in the presence of vancomycin, determined through whole-genome sequencing on the adapted populations. In total, four mutations (in three genes) were identified in uracil-adapted populations, three mutations (in two genes) in L-Valine-adapted populations, and seven mutations (in five genes) in vancomycin-adapted populations (indicated by radial lines in lavender in Figure 7A-C). The mutations’ high frequency and condition-specificity are indicative of their adaptive nature. Furthermore, in the nutrient (uracil and L-Valine) adapted populations the mutated genes are involved in the metabolic pathways of the depleted nutrient (Supplementary Table 3). Additionally, in the vancomycin adapted populations, mutated genes are involved in capsule metabolism (SP_0350/*cps4E*), cell division/cell-wall synthesis (SP_/1067*ftsW*), stringent response (SP_1645/*relA*), membrane transport (SP_1796), and carbohydrate metabolism (SP_2107/*malM*). Although few cases of vancomycin resistance/tolerance have been reported in *S. pneumoniae*, the capsule influences sensitivity to this antibiotic [24, 32, 33], while reduced sensitivity to vancomycin has been reported in *relA* mutants of other Gram-Positive cocci, including *Enterococcus faecalis* [34], vancomycin-resistant *E.faecium* [35], *Staphylococcus aeurus* [36], and cell wall modifications (e.g. thickening) are common features for vancomycin resistance [37, 38].

**Figure 7.**
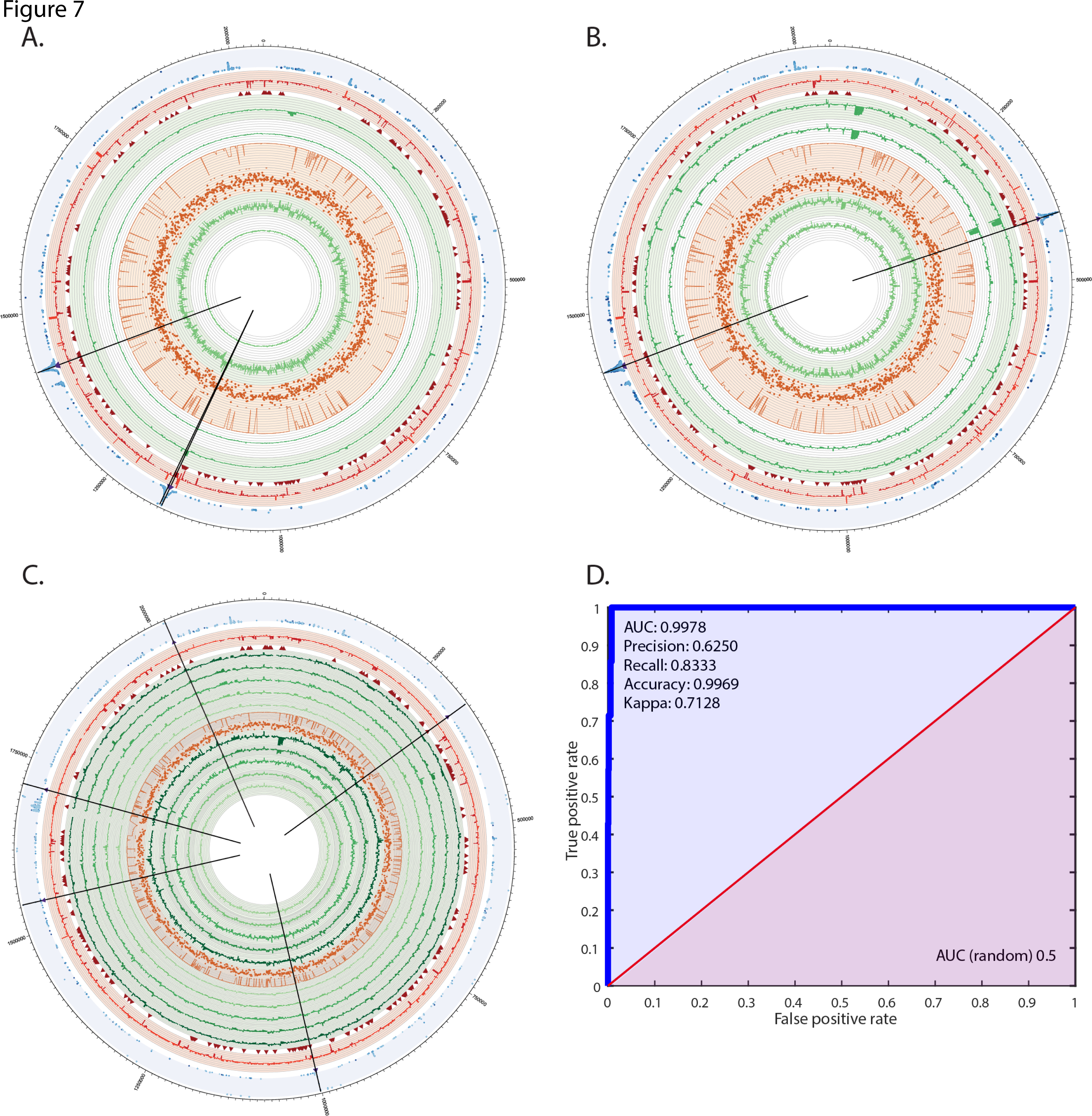
Prediction of adaptive evolution through the integration of machine learning, genome-wide profiles, network characteristics and pan-genome sequence conservation. Pan-genome-wide sequence conservation, RNA-Seq, Tn-Seq and adaptation data are assembled for the uracil (**A.**),L-Valine (**B.**) and vancomycin (**C.**) experiments and visualized by circular plots: 1) Green bar plots represent expression change of parental (the innermost circles) and adapted strains (outside the orange trace); each circle represents a time-point. 2) The orange scatter plot indicates sequence conservation score, while the orange trace is a count of strains that share a gene; 3) Red arrows mark essential genes; 4) Red bar plot represents Tn-Seq fitness change; 5) Blue scatter plot indicates the mutation frequencies, with adapted genes marked by purple arrows and black lines. **D.** Receiver-operator curve (ROC) for SVM classifier. An SVM is trained to distinguish adapted genes from non-adapted genes with high accuracy. Cohen's kappa, precision, recall, and AUROC are reported.

Genotypes of the mutated genes were compared to their homologous genes in 371 *S. pneumoniae* strains that cover the variation present in the pan-genome [39]. Interestingly, the adaptive mutations that arose in the nutrient experiments always resulted in the acquisition of the nutrient-insensitive T4 genotype at these loci (Supplementary Table 3), which is also the shared genotype among the majority of the pan-genome strains, indicative of most strains being tolerant to nutrient deprivation of uracil and L-Valine. In contrast, adaptation to vancomycin results in the acquisition of novel genotypes; i.e. none of the 371 strains carry any of the aT4-vanc mutations, indicative of the fact that very few vancomycin-resistant/tolerant clinical strains have been reported for *S. pneumoniae*. Despite this difference between nutrient and antibiotic adaptive patterns, there are common features to all mutations from all three conditions. For instance, they appear in highly conserved genes, i.e. core genes with high sequence similarity. Next, adaptation data was overlaid with genome-wide profiles and sequence conservation data (Figure 7A-C) in order to visually inspect whether adapted genes overlap with drastic phenotypic changes, transcriptional changes and/or sequence conservation. For example, *carA* (SP_1275) is an adapted gene in the uracil adaptation experiment and it also has both transcriptional and phenotypic importance. While this is a suggestive pattern, at a genome-wide level it is hard to detect such consistent patterns across all three experiments that could be indicative of other likely candidates for adaptive evolution (Figure 7A-C).

To generate a classifier that is able to separate adapted genes (AGs) from non-adapted genes by uncovering hidden patterns in our data, an SVM was built on the Tn-Seq and RNA-Seq profiles, the network characteristics as well as the species-wide sequence conservation data. Importantly, the latter datatype is included because sequence conservation is indicative of genomic plasticity, i. e. it gives insight into the genomic regions that change the most/least and thereby potentially influences the adaptability of each gene. Subsequently, the SVM was trained on the aggregation of all adaptation experiments (uracil, L-Valine and vancomycin), with oversampling of the AGs (see Methods). A total of 1409 data points and 18 features were used, with 10-fold cross-validation and no parameter tuning. In total, 5 out of 6 adapted genes that are on our network are successfully identified as adapted with 3 false positives and 1 false negative (Supplementary Table 3). In cases where one class dominates the dataset (e.g. here we have >99% non-AGs) a high accuracy can even be achieved by a naïve classifier that only selects the more numerous class. Therefore, the observed accuracy of the classifier (99.69%) is compared to a naïve classifier, which performs significantly worse (98.91%, Cohen’s kappa=0.7128, p=0). Furthermore, the sensitivity of the SVM (true positive rate: the proportion of true AGs that are correctly identified) is 83.33%, the specificity (true negative rate: the proportion of true non-AGs that are correctly identified) is 99.77% and the classifier achieved an AUROC (Area Under Receiver Operating Characteristic curve, representing the tradeoff between true positive and false positive rates) of 0.9978, which significantly outperforms a random classifier (AUROC=0.5) and thus indicates that AGs are successfully distinguished from non-AGs (Figure 7D). Importantly, this means that adapted genes indeed share certain common features that are not immediately obvious but can be detected using machine learning. Prior studies of adaptive evolution focus on interpreting adaptive mutations only after they have been acquired, and these interpretations are very specific to the selective pressure under which adaptation has happened in a particular study [40-42]. Instead, the classifier presented here can make *a priori* predictions on which genes will adapt under stress/selective pressure, regardless of the nature of this stress. Thus, we demonstrate that incorporation of different data-types reveals that deterministic factors exist that shape adaptive evolution thereby making it predictable.

## CONCLUSIONS

An important goal here is to determine what type of data is needed to predict a bacterium’s chances of surviving in its environment. We show that a comparison of transcriptional responses between a stress-sensitive and insensitive strain by itself shows stark differences in the number, magnitude and functional tags that are involved in responding to the environment, which are suggestive for their differences in survival-success. The full response thus carries important information; however, a more granular analysis of the response is no less interesting. For instance, both T4 and D39 respond very similarly early on to uracil depletion by ‘appropriately’ upregulating expression of the UMP-pathway, and while T4 maintains a similar response over time, D39’s response is overwhelmed by genome-wide differential expression, resulting in chaos. In addition, components of the stringent response (which is not understood in detail in *S. pneumoniae*) such as genes involved in purine biosynthesis (SP_0044-0056) are down-regulated in both T4 and D39 under amino acid depletion (L-Valine and Glycine; Supplementary file 3). While this shows that particular response mechanisms are activated under stress, it turns out that this is only a partial view. We show that by extending our focus and by taking the temporal genome-wide response into account, it is possible to paint a global and detailed picture of how the organism senses and processes stress. Moreover, we showed previously that it is important to interrogate a bacterial response at both the transcriptional and phenotypic level to uncover network patterns [9], and also here we find that PIGs are critical in enhancing our network coordination analyses, especially when there are a few TIGs (Supplementary File 4). Overall our strategy demonstrates that by integrating temporal transcriptional and phenotypic changes into strain-specific networks, distinct patterns emerge that can be depicted as trajectories in feature space. These temporal trajectories are composed of three types of measurements, Connectedness, Closeness and Representation (CC, CN, RE) that capture the defining network characteristics of a response and objectively quantify a strain’s response into a degree of coordination that reflects survival success. Importantly, we show that the degree of coordination is an evolvable trait; when strains evolve the ability to grow in the absence of a nutrient, or when antibiotic resistance emerges, the network is rewired, increasing coordination and unfolding a focused and targeted response. In other words, selective pressure optimizes a strain’s network coordination, which in turn increases survival success; explaining why network coordination can be used to predict short-term survival outcome. Past adaptation and future adaptive outcome are thereby intricately linked, leading to the possibility of predicting where innovation (adaptation) in the network is most likely to occur. Indeed, we show that by developing a support vector machine that incorporates a wide array of data-types, genes that adapt can be distinguished from those that do not. This indicates that with the right information, adaptation becomes a predictable process.

To improve on the short-term survival outcome and long-term adaptive outcome predictions, it is likely that additional types of data as well as genome-wide networks will be beneficial. For instance, epistatic and regulatory interactions have been shown to influence adaptive evolution [43-46]. It is also possible to include information pertaining to the external environment that the pathogen experiences into a predictive framework. The simultaneous transcriptomic profiling of the host via dual RNA-Seq [47] and cytokine profiling (e.g. determining the state of the host response can allow us to infer the magnitude of host-associated stress the pathogen is experiencing) could also be informative and is something we are currently exploring. Along with the host-response, the infection-causing pathogen potentially experiences competition or participates in cooperation with the resident microbiota of the infection site, which can influence the effectiveness of a given antimicrobial treatment [48]. Therefore, metagenomic profiling of the microbiota from the site of infection may also aid in predicting the survival of a specific pathogen.

To conclude, we demonstrate that network analyses and machine learning make short-term survival outcome and long-term adaptive outcome predictable. Most importantly, the approach is generalizable with respect to the applicability to Gram-positive and Gram-negative bacteria, the emergence of antibiotic resistance, and the applicability to *in vivo* host infection. Thus, our approach offers a primary gateway towards the development of highly accurate infectious disease prognostics.

## MATERIALS AND METHODS

### Bacterial strains, culture media and growth curve assays

*S. pneumoniae* strain TIGR4 (T4; NC_003028.3) is a serotype 4 strain originally isolated from a Norwegian patient [49, 50], Taiwan-19F (19F; NC_012469.1) is a multi-drug resistant strain [51, 52] and D39 (NC_008533) is a commonly used serotype 2 strain originally isolated from a patient about 90 years ago [53]. All gene numbers refer to the T4 genome. Correspondence between homologous genes among the three strains and gene function annotations are described in Supplementary File 3. Unless otherwise specified, *S. pneumoniae* strains were cultivated in Todd Hewitt medium with 5% yeast extract (THY) with 5uL/mL oxyrase (Oxyrase, Inc) or on sheep’s blood agar plates (Northeastern Laboratories) at 37^o^C with 5% CO2. Tn-Seq and RNA-Seq experiments under nutrient-depletion and vancomycin conditions were performed in chemically defined medium (CDM; [9]) and semi-defined minimal medium (SDMM; [21]), respectively. Single strain growth assays were performed at least three times using 96-well plates by taking OD600 measurements on a Tecan Infinite 200 PRO plate reader.

### Tn-Seq experiments, sample preparation and analysis

Six independent transposon libraries were constructed in T4 using transposon Magellan 6 as previously described [10, 11, 21]. Tn-Seq experiments under single nutrient depletion conditions were performed in CDM in the presence or absence of one of the three nutrients: Glycine, uracil and L-Valine. Vancomycin Tn-Seq experiment were performed in SDMM in the presence or absence of 0.1ug/mL vancomycin (MP Biomedicals).

Library preparation, Illumina sequencing, data processing and fitness calculations (*Wi*; representing the growth rate) were performed as previously described [10, 11, 21]. Genes with significant fitness change must satisfy three criteria: 1) Fitness of a gene must be calculated from at least three insertion mutants in both control and experimental conditions. 2) A gene must have a fitness difference greater than 15% (|W_Control_-W_Experimental_|>0.15). 3) W_Control_ and W_Experimental_ must significantly differ in a one sample t-test with Bonferroni correction for multiple testing.

### Temporal RNA-Seq sample collection, preparation and analysis

In nutrient RNA-Seq experiments, T4, D39 and adapted D39 were collected at 30 and 90min after depletion of D39-essential nutrients (Supplementary Table 1). In vancomycin RNA-Seq experiment, T4 and adapted T4 were collected at 10, 20, 30, 45, 60 and 90min post-vancomycin (1x MIC) treatment. Cell pellets were collected by centrifugation at 4000 rpm at 4^o^C and snap frozen and stored at -80^o^C until RNA isolation by RNeasy Mini Kit (Qiagen). 400ng of total RNA from each sample was used for generating cDNA libraries following the RNAtag-Seq protocol [54] as previously described [9]. PCR amplified cDNA libraries were sequenced on an Illumina NextSeq500 generating a high sequencing depth of ~7.5 million reads per sample [55]. RNA-Seq data was analyzed using an in-house developed analysis pipeline. In brief, raw reads are demultiplexed by 5’ and 3’ indices [54], trimmed to 59 base pairs, and quality filtered (96% sequence quality>Q14). Filtered reads are mapped to the corresponding reference genomes using bowtie2 with the --very-sensitive option (-D 20 –R 3 –N 0 –L 20 –i S, 1, 0.50) [56]. Mapped reads are aggregated by featureCount and differential expression is calculated with DESeq2 [57, 58]. In each pair-wise differential expression comparison, significant differential expression is filtered based on two criteria: |log2foldchange| > 1 and adjusted p-value (padj) <0.05. All differential expression comparisons are made between the presence and absence of the nutrient at the same time point.

### Experimental evolution and whole-genome sequencing

D39 and T4 were used as parental strains in nutrient-depletion and vancomycin evolution experiments, respectively. Four replicate populations were grown in fresh CDM with decreasing concentration of uracil or L-Val for nutrient adaptation populations, or increasing concentration of vancomycin for antibiotic adaptation populations. Four replicate populations were serial passaged in CDM as controls for background adaptations in nutrient adaptation experiments. When populations had adapted a single colony was picked from each experiment, checked for its adaptive phenotype by growth curve experiments. Genomic DNA was isolated from adapted populations and single strains using a DNase Blood and Tissue kit (Qiagen), concentrations of genomic DNA were measured on a Qubit 3.0 fluorometer (Invitrogen) and diluted to 5ng/uL for library preparation using a Nextera kit (Illumina). Libraries were sequenced on an Illumina NextSeq500 and reads were mapped to their corresponding reference genomes. Mutations were identified using the breseq pipeline with polymorphism mode for populations and consensus mode for adapted strains [59]. Adaptive mutations in each experiment are determined based on the following criteria: 1) mutation frequency is greater than 50% in at least one replicate population, and 2) this mutation is not present in any CDM-background adapted populations and 3)the mutation is a nonsense or missense mutation.

### Determination of relative minimal inhibitory concentration (MIC) by microdilution

1 to 5 x 10^5^ CFU of mid-exponential T4 in 100uL was diluted with 100uL of fresh medium with vancomycin to achieve a gradient of final concentrations from 0 to 0.5ug/mL in 96-well plates. Each concentration was tested in triplicates. Growth was monitored on a Tecan Infinite 200 PRO plate reader at 37^o^C for 16 hours. MIC is determined as the lowest concentration that abolishes bacterial growth (Supplementary Figure 1C).

### Fluorescent microscopy

Wild-type and vancomycin adapted T4 were grown to mid-exponential phase. Half of the culture was left untreated, while the other half was exposed to 0.24ug/mL of vancomycin for 60 minutes. 1×10^8^ CFUs were collected by centrifugation, resuspended in 20uL of PBS and stained with Syto9 (DNA stain) and FM4-64 (cell membrane stain) for 10 minutes at room temperature. 1uL of stained cells were imaged on an Olympus IX83 microscope system with an ORCA-Flash4.0 camera (Hamamatsu) and a 60x oil immersion objective. Phase contrast and fluorescence images through GFP and RFP channels were taken for each sample. Microscopy of each sample was repeated with at least three technical replicates. Images were modified for publication using Fiji [60]. Cell numbers per chain was visually quantified based on 1000 *S. pneumoniae* chains from each treatment group using at least three technical replicate micrographs.

### Strain-specific metabolic model construction

Thirty-six reactions were manually added to the previously described T4 model [9] using the COBRA toolbox based on updated information from three databases (NCBI, KEGG and BiGG) and literature [61, 62]. Metabolite and reaction IDs were cross-referenced to follow the BiGG naming convention [63]. Gene-reaction associations in the updated T4 metabolic model were adjusted into three strain-specific models based on the correspondence table (Supplementary Table 1). For visualization of the metabolic models, a map of the *S. pneumoniae* metabolism was constructed using Escher [64] referencing the KEGG pathway base (Supplementary Figure 2).

### Spectral Clustering of the *S. pneumoniae* pan-genome

Complete, annotated genomes from 22 reference strains (RefSeq 58) and contigs from 350 clinical strains [39] were assembled for data analysis. The contigs were annotated using the PATRIC Genome annotation service to identify coding sequences [65]. A total of 820,754 amino acid sequences from the 372 strains were assembled. In order to reduce redundancy and expedite clustering, representative sequences were selected using a boundary forest algorithm [66], with Smith-Waterman distance as the similarity measure. This decreased the number of sequences to 17,000 representatives. Pairwise distances between representatives were computed to generate a sequence similarity matrix (S). The gaussian kernel of S was thresholded and transformed to an adjacency matrix. Spectral clustering with normalization of the Laplacian was performed to generate sequence clusters [67]. Since we had no prior knowledge of what the most appropriate number of clusters would be, we scanned the range of 1000 to 10,000 clusters, and computed the sum of squared errors (SSE) on all clusters, for each cluster set. SSE was minimized at 4300 clusters, therefore, we determined this to be the appropriate number of clusters of homologous genes in the *S. pneumoniae* pan-genome. Sequences in each gene cluster were aligned using Clustal Omega [68], and the average pairwise Smith-Waterman distance within each cluster was computed. In the case of large clusters (containing >50 sequences), 50 random sequences were selected for pairwise distance calculation. We define gene conservation as -log(mean(distance)) within a cluster, and count (number of strains that share the gene) of sequences in each cluster.

### Network coordination analysis

We define 3 criteria for metabolic coordination: connectedness (CC), closeness (CN) and representation (RE) in the metabolic network. Number of connected components (NCC) is used as the metric for connectedness. For each experiment, connected components were determined using the components function in the igraph package [69]. Since the expected NCC heavily depends on the number of nodes selected, and the network architecture, in order to test whether the observed NCC is significantly lower than expected, we apply permutation testing on random selection of nodes on the network as follows: In an experiment with M responsive genes on the network, we generate 1000 sets of M random genes, and compute the NCC for each permutation. The empirical one-tailed p-value for this experiment is the proportion of permutations in which we observed fewer NCC than the responsive genes in the experiment. A response is connected if the empirical p-value for the NCC permutation testing is <0.01. To determine closeness of responsive genes, the average length of shortest paths is computed for each pair of genes. Since biological pathways may appear as long chains with few branches, it is possible to have a connected component of TIGs and/or PIGs arranged in a line, with a high average pairwise distance. In order to avoid such skew, we considered any responsive gene pairs that appear in the same component to be at distance 0 to each other by assigning each edge on the network a weight of 0 if it connects two responsive genes, and 1 otherwise. If there is no path connecting the two components, the distance between this pair is replaced by the diameter of the network+1 (i.e. 21 in our network), to avoid infinite values. Similar to connectedness evaluation, permutation testing is applied to the average network distance. A response is “close” if the empirical p-value for the distance permutation testing is <0.01. To assess whether TIGs and PIGs were significantly highly represented in the metabolic network we consider N, the total number of responsive genes, and M, the subset of N that appear on the network. The probability of observing M or more genes on the network, given N total responsive genes in the genome (p(m>M|N)) is computed assuming a hypergeometric distribution. A response is metabolically well-represented if this probability is <0.01.

### Support Vector Machine Classification of Adapted Genes

A support vector machine (SVM) using a gaussian kernel is trained and cross-validated using the fitcsvm function in MATLAB to distinguish whether a gene will contain adaptive mutations or not. The model was trained on network parameters (degree, transitivity, centrality), TnSeq, RNAseq and sequence conservation (count, or number of occurrences across the pan-genome, and sequence similarity) of each gene. Data from the dependent (parental) strains from the uracil (D39), L-Valine (D39) and vancomycin (T4) experiments were assembled into a set of 1283 data points with 18 features that were standardized. Genes that were not represented on the metabolic network were excluded. Each observation was then labeled as AG or non-AG. Because the number of AGs is very small (6 out of 1283), we applied synthetic minority oversampling [70] until 10% of the observations were AGs. The SVM was trained on a total of 1409 data points (1283 experimental and 126 synthetic) using 10-fold cross-validation, and report the average accuracy, kappa, precision and recall on the 10 cross-validation sets.

### Statistical analysis

Quantification and statistical analysis are described in the above Method Details section, Supplementary Table2 and in figure legends (Figures 2, 3, 5, S4).

## List of abbreviations

AG: adapted gene
AUROC: area under receiver operating characteristic curve
CC: connectedness
CDM: chemically defined medium
CFU: colony forming unit
CN: closeness
MIC: minimum inhibitory concentration
NCC: number of connected components
PIG: phenotypically important gene
RE: representation
RNA-Seq: RNA-Sequencing
SDMM: semi-defined minimal medium
SSE: sum of squared errors
SVM: support vector machine
TIG: transcriptionally important gene
Tn-Seq: transposon insertion sequencing
UMP: uridine monophosphate

## Accession numbers

The datasets generated during the current study are available as Supplementary Files and in the Sequence Read Archive (SRX2039176, SRX2039177, SRP156493 and SRP156489).

## Funding

This work was supported by NIH R01 AI110724 and U01 AI124302.

## Authors’ contributions

TvO devised the study, ZZ performed the wet-lab experiments, DS performed the network analyses, clustering and machine learning. JB consulted on the clustering and machine learning. AP constructed the metabolic maps. ZZ, DS and TvO analyzed data, interpreted results and wrote the manuscript. ZZ, DS, AP, JB and TvO approved the final manuscript.

## Acknowledgements

DNA sequencing was performed at the Boston College Sequencing Core. The authors wish to thank Jon Anthony for running the Aerobio sequencing analyses pipeline.

## Conflict of interest

The authors declare that they have no conflict of interest.

---

**Supplementary File 1:** Supplementary information

**Supplementary File 2:** iSP16 consensus model.

**Supplementary File 3:** Tn-Seq and temporal RNA-Seq data in this study.

**Supplementary File 4:** Network analysis with TIGs and PIGs, and only TIGs.

